# Neurodevelopmental wiring deficits in the Ts65Dn mouse model of Down syndrome

**DOI:** 10.1101/784314

**Authors:** Shruti Jain, Christina A. Watts, Wilson C.J. Chung, Kristy Welshhans

## Abstract

Down syndrome is the most common genetic cause of intellectual disability and occurs due to the trisomy of human chromosome 21. Adolescent and adult brains from humans with Down syndrome exhibit various neurological phenotypes including a reduction in the size of the corpus callosum, hippocampal commissure and anterior commissure. However, it is unclear when and how these interhemispheric connectivity defects arise. Using the Ts65Dn mouse model of Down syndrome, we examined interhemispheric connectivity in postnatal day 0 (P0) Ts65Dn mouse brains. We find that there is no change in the volume of the corpus callosum or anterior commissure in P0 Ts65Dn mice. However, the volume of the hippocampal commissure is significantly reduced in P0 Ts65Dn mice, and this may contribute to the impaired learning and memory phenotype of this disorder. Interhemispheric connectivity defects that arise during development may be due to disrupted axon growth. In line with this, we find that developing hippocampal neurons display reduced axon length *in vitro*, as compared to neurons from their euploid littermates. This study is the first to report the presence of defective interhemispheric connectivity at the time of birth in Ts65Dn mice, providing evidence that early therapeutic intervention may be an effective time window for the treatment of Down syndrome.

## 1. Introduction

Down syndrome results from the trisomy of human chromosome 21. Furthermore, it is the most common genetic cause of intellectual disability and the most prevalent congenital disorder, occurring in about 1 of every 792 live births [1]. Studies have shown that partial and complete trisomy of human chromosome 21 (HSA21) leads to various neurological phenotypes in Down syndrome, including reduced brain weight and volume [2, 3] and a decreased number of proliferating neurons during the fetal period [4]. Adolescents and adults with Down syndrome have a reduction in the size of the corpus callosum, hippocampus [5, 6] and anterior commissure [7]. Multiple studies have identified neurological abnormalities in Down syndrome, which may contribute to the intellectual disability phenotype of this disorder; however the molecular mechanisms underlying these phenotypes remain unclear.

The most commonly used mouse model of Down syndrome is the Ts65Dn strain [8, 9]. These mice have over 100 genes triplicated that are orthologs to HSA21 plus about 30 protein coding genes that derive from mouse chromosome 17 and are not orthologs of HSA21 genes [10, 11]. Ts65Dn mice also replicate many of the behavioral phenotypes of Down syndrome [9, 12, 13]. Studies have identified neurological changes that occur during development and may contribute to these phenotypes. For example, the pyramidal cell layer and wall of the CA1 hippocampus is smaller in Ts65Dn mice during embryonic development, and there is a reduction in synaptogenesis during the early postnatal period [14]. In line with this finding, there is a decrease in proliferation in multiple hippocampal regions of Down syndrome fetuses and the dentate gyrus of postnatal day 2 (P2) Ts65Dn mice [4, 15, 16]. At P6 in the Ts65Dn mouse hippocampus, there is still a decrease in the number of granule cells in the dentate gyrus, but the number of cells in the pyramidal cell layer is not different from controls [17]. More recent studies have demonstrated that treatment of newborn Ts65Dn mice with a sonic hedgehog agonist can rescue proliferation deficits in the cerebellum, but not the dentate gyrus [18]; however prenatal treatment of mice with fluoxetine can rescue proliferation defects in the hippocampus [16]. Interestingly, both of these treatments improve performance of Ts65Dn mice on hippocampal-dependent tasks [16, 18]. However, we do not know how changes in the formation of connectivity during development may also contribute to this disorder. Understanding changes in neural wiring during development, as well as the cellular and molecular mechanisms contributing to these changes, is essential to further our understanding of Down syndrome and may provide insight into treatments for this disorder.

Here we employ Ts65Dn mice, the most widely used mouse model of Down syndrome, to examine which connectivity deficits are present at birth. We find that the volume of the hippocampal commissure is significantly reduced in postnatal day 0 (P0) Ts65Dn mice. This is likely due to deficits in axon growth because hippocampal neurons from Ts65Dn mice have significantly decreased axon length *in vitro*. Interestingly, we find that the volume of the corpus callosum and anterior commissure are not altered in P0 Ts65Dn mice. Taken together, these data suggest that deficits in the formation of long-distance connectivity during development in specific brain regions may contribute to the intellectual disability phenotype of Down syndrome.

## 2. Materials and Methods

### 2.1 Animals and cell culture

All experimental procedures were approved by the Institutional Animal Care and Use Committee at Kent State University. Ts65Dn (B6EiC3Sn.BLiA-Ts(17^16^)65Dn/DnJ) and B6EiC3Sn.BLiAF1/J mice were obtained from The Jackson Laboratory. Ts65Dn mice were obtained by crossing Ts65Dn trisomic female mice to B6EiC3Sn.BLiAF1/J male mice. Brains from postnatal day 0 (P0) animals were collected within 24 hours of birth. Each experiment was carried out using equal numbers of wild type and trisomic mice from at least three different litters. Each experiment had litters from at least three independent dam/sire pairs. Tail clips from animals were collected and genotyped using breakpoint PCR as described previously [19]. All pups were also sexed using SRY primers (forward: TTG TCT AGA GAG CAT GGA GGG CCA TGT CAA; reverse: CCA CTC CTC TGT GAC ACT TTA GCC CTC CGA), however no significant differences were seen when the data were compared between males and females. Therefore, wild type and trisomic groups contain mice of both sexes.

Hippocampal cultures from trisomic Ts65Dn animals (Ts65Dn) and their wild-type euploid littermates (WT) were prepared in line with previously published techniques [20-22]. Briefly, hippocampi were transferred to 0.25% trypsin and incubated at 37°C for 5 minutes. Hippocampi were then rinsed twice in prewarmed Hank’s Balanced Salt Solution (HBSS) without calcium, magnesium and phenol red (Corning) at 37°C for 5 minutes each rinse. Next, hippocampi were transferred to 1 ml of Minimum Essential Medium (MEM) (Corning) and mechanically dissociated. Cells were counted and 15,000 cells were plated on each poly-l-lysine coated (Sigma) glass coverslip (Carolina Biological) in MEM with FBS. Two hours after plating, the media was replaced with fresh pre-warmed Neurobasal (Gibco) with 2% B27 and 1X glutamax.

### 2.2 Axon outgrowth and branching

Axon outgrowth and branching experiments were performed on P0 Ts65Dn and WT hippocampal neuronal cultures. Hippocampal neurons were grown for 2 DIV, and then fixed and immunostained with mouse anti-β-tubulin (1:1000; DSHB) as previously described [20]. The following secondary antibody was used: goat anti-mouse Alexa 488 (Jackson ImmunoResearch). Coverslips were briefly rinsed with deionized water and mounted using Prolong Gold antifade mounting media (Life Technologies). Axon outgrowth and branching was quantified as previously described [20], and explained in the analysis section below.

### 2.3 Immunohistochemistry

For *in vivo* immunohistochemistry (IHC) experiments, P0 brains from Ts65Dn and WT animals were fixed in 4% PFA for 6 hours. Brains were then transferred to 30% sucrose/1X phosphate buffer saline (PBS) solution and allowed to sink to the bottom of the vial overnight. Brains were stored at 4° C until use. Prior to sectioning, brains were frozen in OCT (Fisher Healthcare) and coronal sections of 25µm thickness were obtained using a cryostat (Leica CM 1950). Sections were mounted to gelatin coated glass slides (Sigma Aldrich).

To perform IHC, sections were first rinsed 3 × 5 minutes with 1X tris buffer saline (TBS; pH 7.6), and then incubated in 1% H_2_O_2_/1X TBS solution for 20 minutes. Next, sections were washed 2 × 5 minutes in 1X TBS for five minutes, followed by incubation in 1X TBS/ 0.3% Triton-X-100 (1X TBS-T). Sections were then incubated with rat anti-L1 antibody (1:500; Milipore) in 2% normal goat serum and 1X TBS-T for 48 hours at 4° C in the dark. Following primary antibody incubation, the sections were washed 3 × 5 minutes with 1X TBS-T, followed by application of biotinylated goat anti-rat antibody (1:500; Vector Laboratories) for two hours. After secondary antibody incubation, sections were again washed 3 × 5 minutes with 1X TBS-T. Sections were then treated with avidin/biotinylated enzyme complex (1:500; Vector Laboratories) for two hours in the dark at room temperature. Sections were washed with 1X TBS 3 × 5 minutes followed by treatment with 0.05% 3,3’-Diaminobenzidine (Sigma Aldrich) and 0.01% H_2_O_2_/1X TBS for 20 minutes. Slides were washed thoroughly with 1X TBS 3 × 5 minutes. Finally, slides were dehydrated with ethanol, cleared with xylene and mounted using DPX (Sigma Aldrich).

### 2.4 Image acquisition and analysis

For image acquisition of axon morphology, an Olympus IX81 microscope, combined with a SensiCam QE CCD camera (Cooke), was used. Only cells with a pyramidal morphology were included in the analysis. Axons and dendrites were initially defined using tau and MAP2 staining. Based on this staining, we found that we can reliably define the axon as the longest neurite on the cell, which extended at least 3 times the length of the next longest neurite.

In experiments examining axon length, ImageJ was used to measure the length of the primary axon (i.e. longest neurite) from the cell body to the center of the axonal growth cone. Axon length does not include the length of branches. In experiments examining axon branching, primary branches were manually counted along the entire length of the axon. A primary branch was characterized as a process greater than 5μm in length extending out from the primary axon. Number of branches was defined as the total number of primary branches on an axon, in line with previous studies [23-25]. Growth cones were defined by enclosing the entire perimeter of the growth cone including central domain, lamellipodia, and filopodia.

For image acquisition of brain sections, an Olympus microscope equipped with a SC-30 Olympus camera was used. Images were taken using an Olympus 4X objective. In order to define a specific region for corpus callosum analysis, we started our analysis with the first anterior section that contained callosal fibers visibly crossing the midline and we continued our analysis until we reached the first section that showed a visible break in midline crossing of callosal fibers. All sections between these two border regions were used for analysis. The distance between each immunostained serial coronal section was 75 µm. Image analysis was performed using NIH ImageJ software. The thickness of the corpus callosum was measured in 3 areas in the coronal sections: (1) the fiber bundle where the arch of the corpus callosum is at its peak in the left hemisphere, (2) the fiber bundle where the arch of the corpus callosum is at its peak in the right hemisphere, and (3) the fiber bundle at the midline of the corpus callosum. To quantify the volume of the corpus callosum, the area encompassing the entire corpus callosum between the peak of the arch in both the left and right hemispheres was calculated, and then multiplied by the coronal thickness between each immunostained section (i.e. 75 µm). To quantify the area of the anterior commissure, the area encompassing the entire anterior commissure was calculated. To quantify the volume of the hippocampal commissure, we outlined the area encompassing the entire hippocampal commissure to calculate the area of this fiber bundle. The calculated area was then multiplied by the coronal section thickness between each stained section (i.e. 75 µm).

Depending on the parametric or non-parametric nature of data, a variety of statistical tests were applied to experimental data using SPSS (IBM) software, including Mann-Whitney, ANOVA, and appropriate post hoc tests. The specific statistical test used in each experiment is given in its figure legend. Significance was set as *p* ≤ 0.05. In all graphs, error bars represent SEM and “n” represents the number of growth cones analyzed for *in vitro* experiments, and the number of animals analyzed for *in vivo* experiments.

## 3. Results

### 3.1 Corpus callosum and anterior commissure volume are not altered in a mouse model of Down syndrome

Here, we employed the most widely used mouse model of Down syndrome (Ts65Dn) to determine if there are *in vivo* connectivity defects during early postnatal development in this disorder. Brains from P0 animals were fixed, sectioned and then immunostained with an L1 antibody to specifically visualize the corpus callosum, anterior commissure, and hippocampal commissure. The total average volume of the corpus callosum was not significantly different between Ts65Dn and wild type (euploid) animals (Figure 1A-E). In addition, there is no significant difference in corpus callosum volume between Ts65Dn and wild type mice when examined by individual section, from anterior to posterior (Figure 1F).

**Figure 1:**
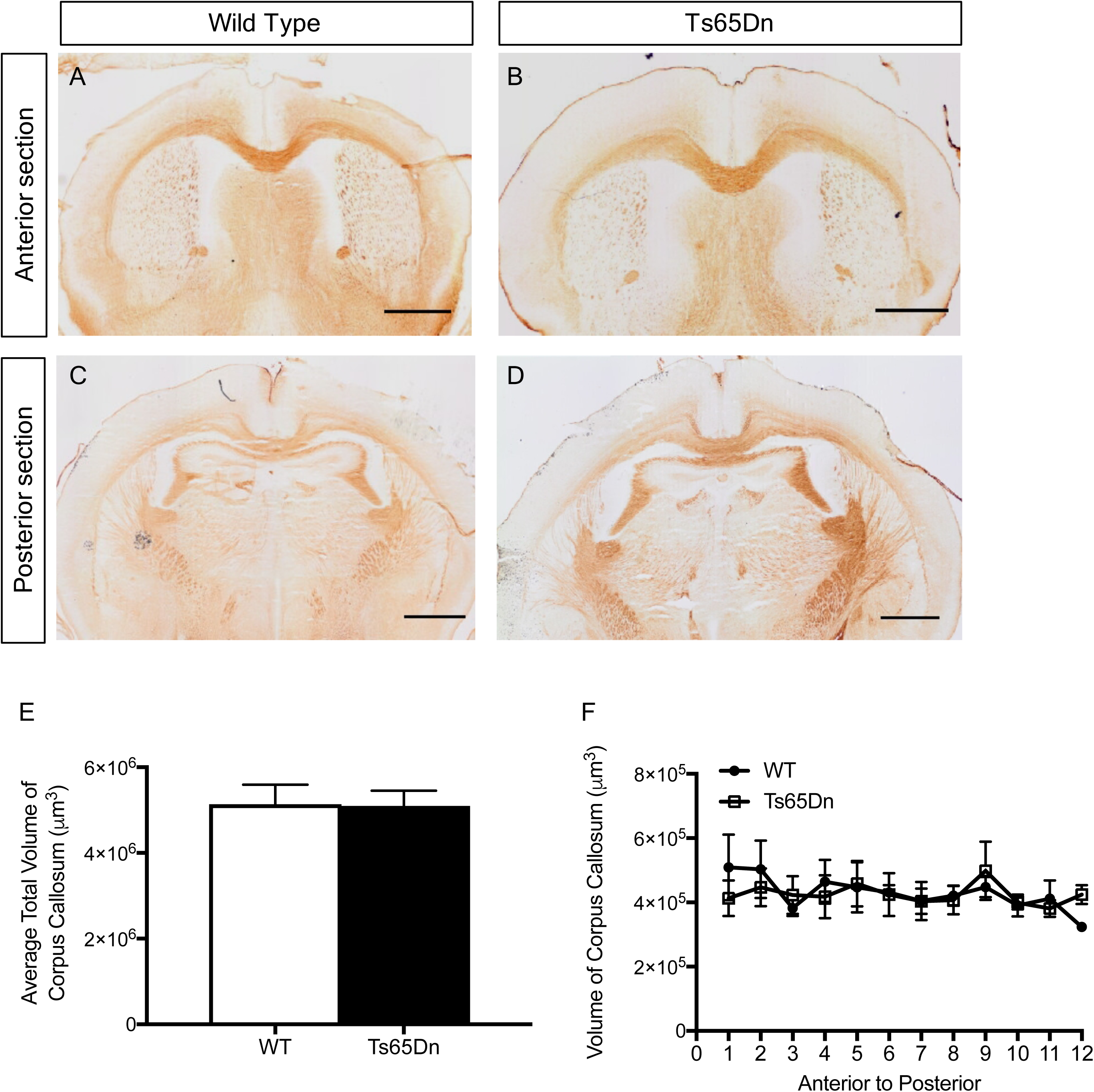
The corpus callosum is not altered in Ts65Dn mice. **A-D**, P0 brains from wild type and Ts65Dn animals were fixed, sectioned and stained for L1. Representative sections from the anterior (**A, B**) and posterior (**C, D**) ends of the corpus callosum used in the analysis are shown. Scale bar, 100µm. **E,** The average total volume of the corpus callosum is not significantly different between P0 Ts65Dn and wild type mice. p=0.95, Student’s t-test. **F,** The volume of the corpus callosum in individual sections is shown, from anterior to posterior. An ANCOVA (between subjects factor: genotype; covariate: section) revealed no main effect of genotype (p=0.882) or section (p=0.065). WT, n=4 (2 male & 2 female mice); Ts65Dn, n=4 (2 male & 2 female mice). These animals derived from 3 separate litters.

Next, we measured corpus callosum thickness in three regions of the fiber bundle: (1) where the arch is at its peak in the left hemisphere, (2) where the arch is at its peak in the right hemisphere, and (3) at the midline of the corpus callosum. No significant differences in the average thickness between Ts65Dn and wild type brains were detected in any of these three regions (Figure 2A). Furthermore, no significant differences were found when we examined these thicknesses in individual sections (Figure 2B-D). Similarly, no significant difference in the average area of the anterior commissure between Ts65Dn and wild type animals was detected (Figure 3D). These results demonstrate that there is no change in the volume of the corpus callosum or anterior commissure in brains of early postnatal Ts65Dn animals. However, most studies have demonstrated that there are connectivity defects in the corpus callosum and anterior commissure in adult humans with Down syndrome [5-7, 26, 27]. Thus, future studies are needed to determine where, when, and how these defects arise.

**Figure 2:**
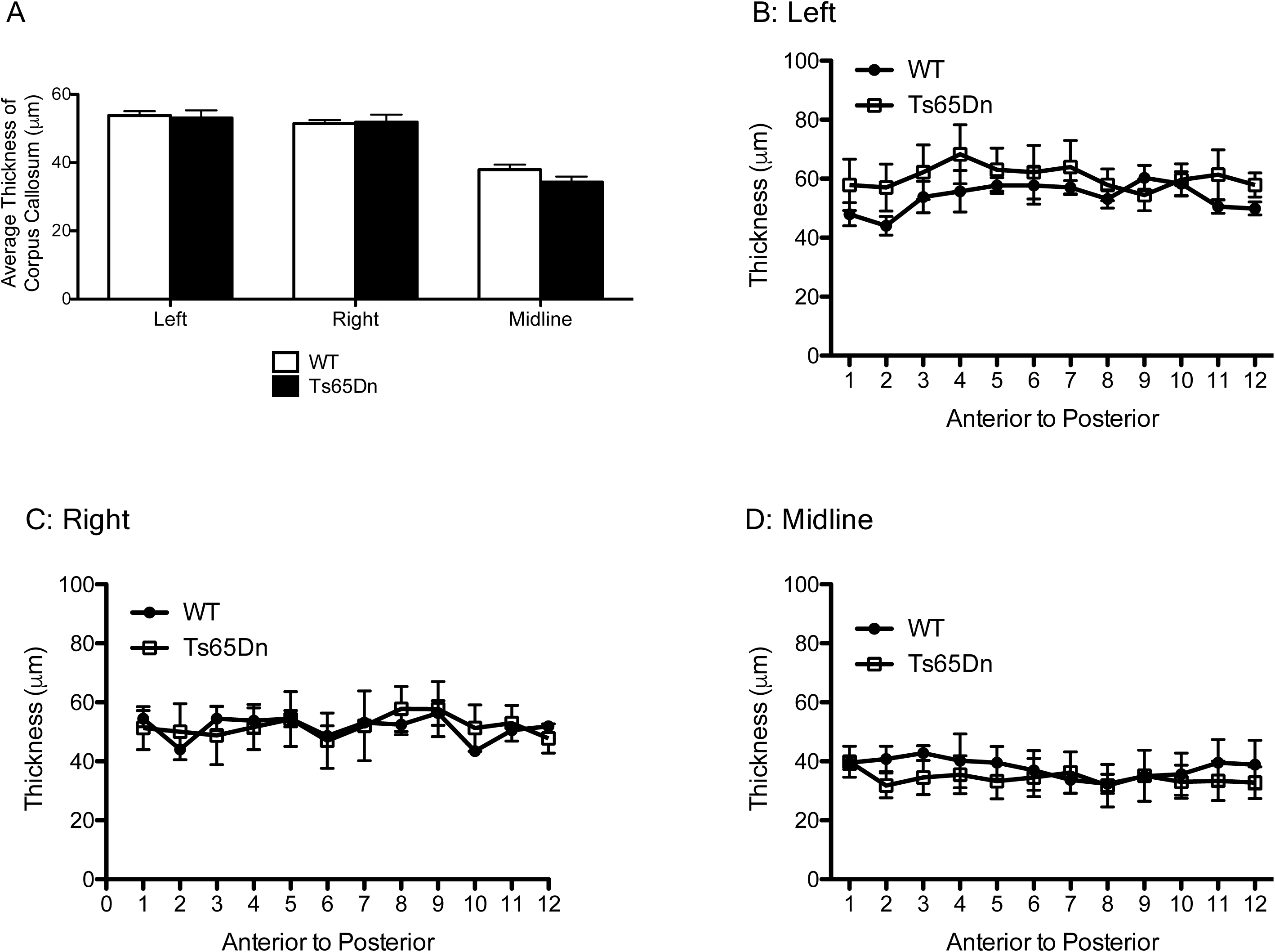
Average thickness of the corpus callosum is not altered in Ts65Dn mice. **A**, The thickness of the corpus callosum was measured in three regions of the fiber bundle: (1) where the arch is at its peak in the left hemisphere, (2) where the arch is at its peak in the right hemisphere, and (3) at the midline of the corpus callosum. Left hemisphere, p=0.977, Mann-Whitney; Right hemisphere, p=0.504 Mann-Whitney; Midline, p=0.333, Student’s t-test. WT, n=4 (2 male & 2 female mice); Ts65Dn, n=4 (2 male & 2 female mice). **B-D**, The thickness of the corpus callosum was examined from anterior to posterior in individual sections. This thickness was measured in the left hemisphere (**B**), right hemisphere (**C**), and midline (**D**). An ANCOVA (between subjects factor: genotype; covariate: section) revealed no main effect of genotype (B, p=0.766; C, p=0.869; D, p=0.110) or section (B, p=0.461; C, p=0.941; D, p=0.287). WT, n=4 (2 male & 2 female mice); Ts65Dn, n=4 (2 male & 2 female mice). These animals derived from 3 separate litters.

**Figure 3:**
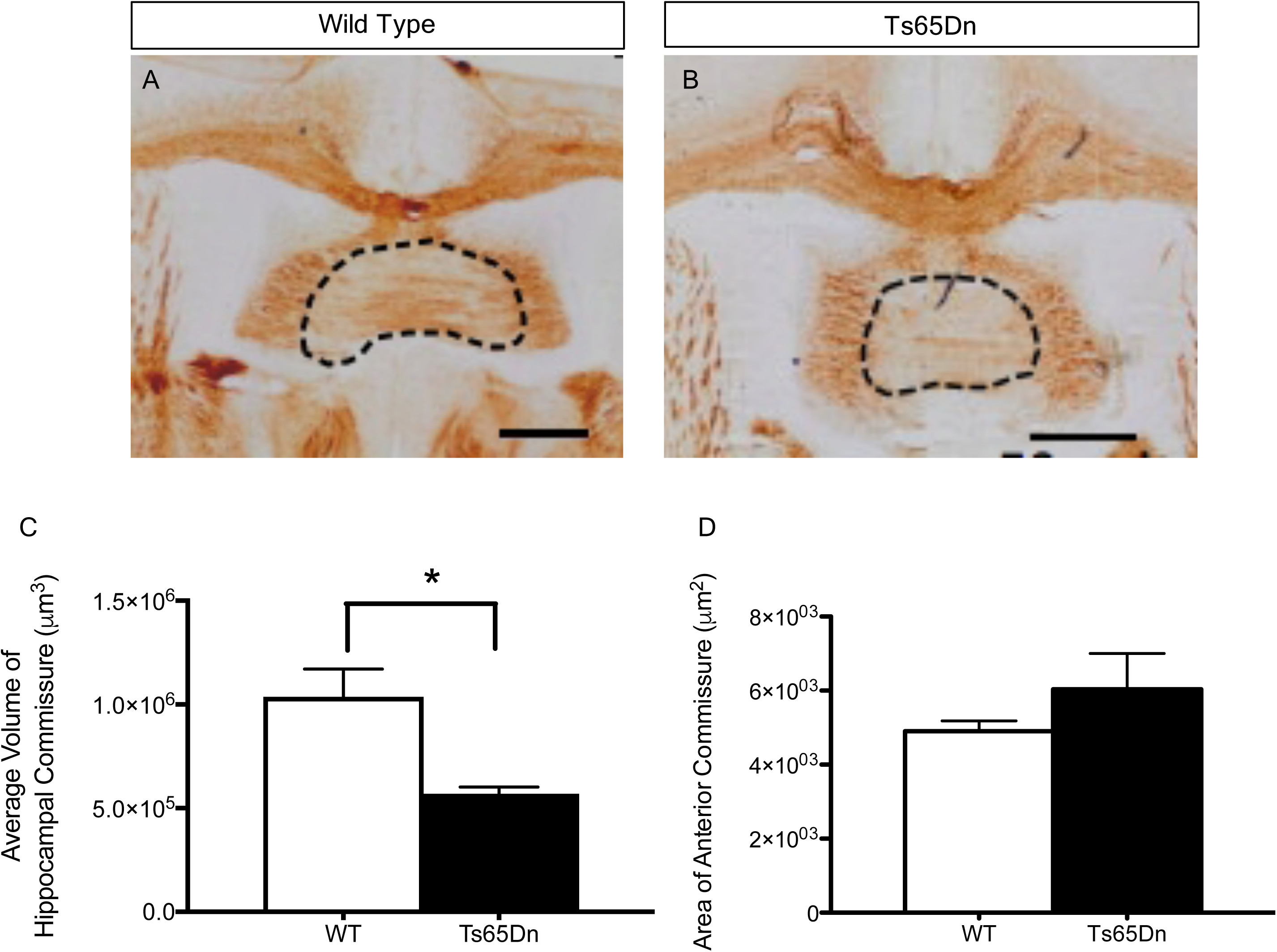
Volume of the hippocampal commissure is significantly smaller in Ts65Dn mice. P0 brains were fixed and sectioned coronally at 25µm thickness. Axonal tracts in brain sections were labeled using L1 immunostaining. Dotted lines outline the hippocampal commissure of WT (**A**) and Ts65Dn mice (**B**) at P0. Scale bar, 50µm. **C**, The average volume of the hippocampal commissure is significantly reduced in Ts65Dn mice as compared to their wild type littermates. WT, n=4 (2 male & 2 female mice); Ts65Dn, n=4 (2 male & 2 female mice). *p ≤ 0.05, Mann-Whitney. **D,** The area of the anterior commissure is not significantly different between Ts65Dn animals and their wild type littermates. WT, n=4 (2 male & 2 female mice); Ts65Dn, n=4 (2 male & 2 female mice). p=0.343, Mann-Whitney. These animals derived from 3 separate litters.

### 3.2 Hippocampal commissure volume is reduced in a mouse model of Down syndrome

We examined the volume of the hippocampal commissure and found that it was significantly reduced in Ts65Dn mice as compared to their wild type littermates (Figure 3A-C). These data suggest that there are defects in hippocampal connectivity formation during development.

### 3.3 Axon outgrowth is reduced in Ts65Dn mice

We next sought to determine if the significant reduction in hippocampal commissure volume observed in Ts65Dn animals may be due to defects in axon growth during development. We compared axon length, number of branches per axon and growth cone area between Ts65Dn neurons and their wild type littermates. Staining of cultured P0 hippocampal neurons for β-tubulin showed that there was a significant reduction in axon length and number of branches per axon in Ts65Dn neurons as compared to their wild type littermates (Figure 4A-D). However, growth cone area was not significantly different between Ts65Dn mice and their wild type littermates (Figure 4E). These data suggest that deficits in axon growth contribute to the hippocampal commissure deficits present in P0 Ts65Dn mice.

**Figure 4:**
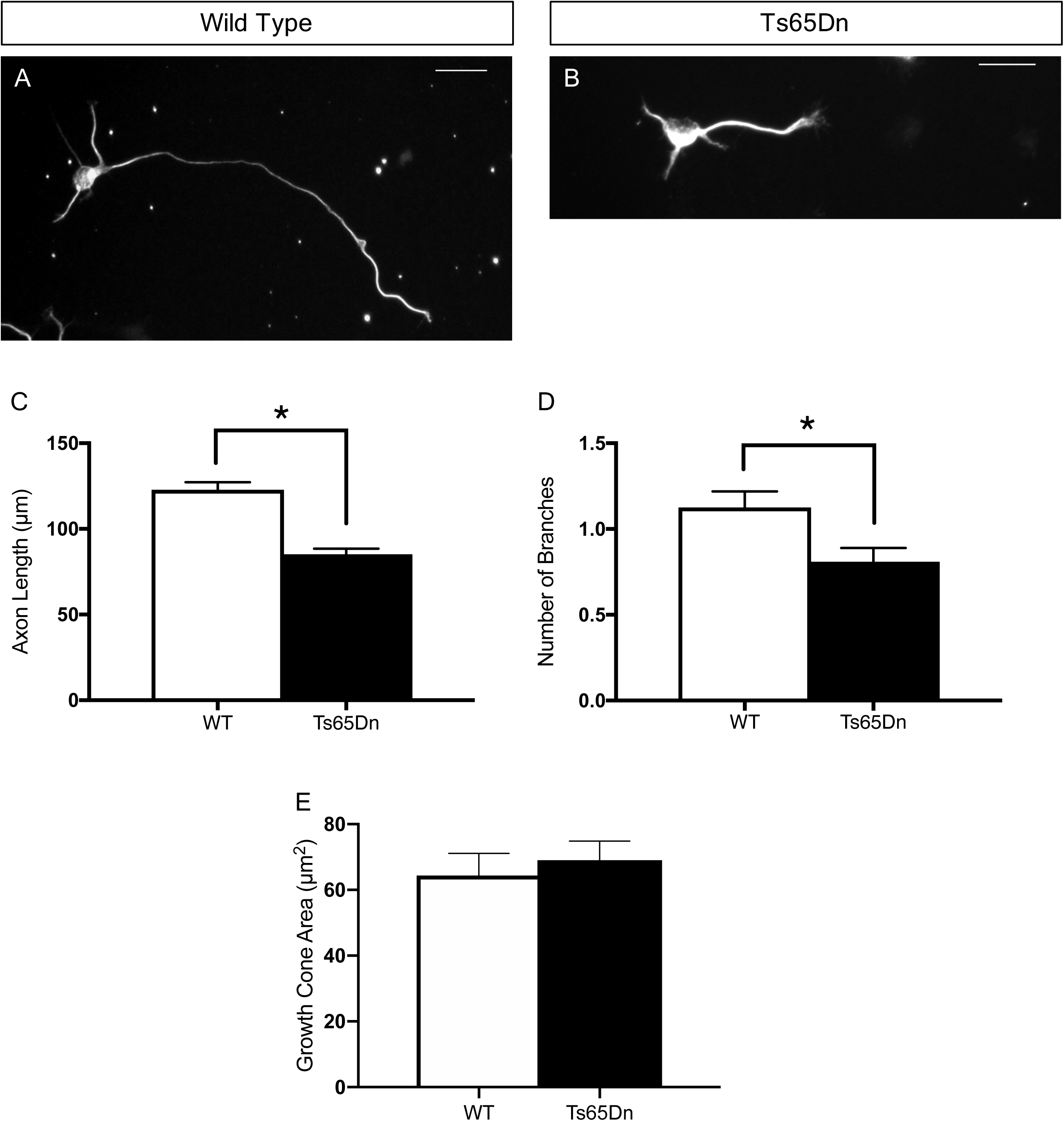
Ts65Dn neurons have reduced axonal outgrowth. **A-C**, Primary hippocampal neurons were grown for 2 DIV and immunostained for β-tubulin. Ts65Dn neurons have significantly shorter axons, as compared to neurons from their wild type littermates. Scale bars, 25μm. *p ≤ 0.05, Mann-Whitney. WT, n=119; Ts65Dn, n=136. **D**, The number of primary branches on Ts65Dn neurons is significant reduced as compared to their wild type littermates. WT, n=119; Ts65Dn, n=136. *p ≤ 0.05, Mann-Whitney. **E,** Growth cone area is not significantly different between Ts65Dn neurons and neurons from their wild type littermates. WT=60; Ts65Dn, n=60. p=0.26, Mann-Whitney.

## 4. Discussion

### 4.1 Deficits in axon growth in Down syndrome

Triplication of genes in the Ts65Dn Down syndrome mouse model leads to stunted axon outgrowth of early postnatal hippocampal neurons (Figure 4C). In contrast to our results, a recent study demonstrated that axon length was increased in hippocampal neurons from Ts65Dn mice [28]. However, these neurons were plated on laminin, whereas in the current study we plated on poly-L-lysine. It is known that substrate and guidance cues directly regulate axon length, and these differences may be magnified by the triplication of genes in this mouse model of Down syndrome. In support of the current findings, two separate studies examining cultured neurons find decreased axon length associated with Down syndrome. These studies employ: (1) cortical neurons from a human fetus with Down syndrome [29], and (2) iPSCs created from individuals with Down syndrome that have been differentiated to neurons [30]. However, future studies will need to further investigate how substrate and guidance cues differentially affect axon growth and guidance in Down syndrome.

The changes in axon length are also in line with our prior research, demonstrating that overexpression of Down syndrome cell adhesion molecule (*Dscam*) results in a severe reduction in axon length and branching [20]. *DSCAM* is located on HSA21, and is not only a cell adhesion molecule, but also a receptor for an axon guidance molecule. DSCAM is highly expressed in both the developing nervous system and adult brain in vertebrates [31-34], and is elevated in adults with Down syndrome [35]. Taken together, these data suggest that increased levels of *Dscam* in Down syndrome may contribute to the axon length phenotype observed in the mouse model of this disorder, however further studies are needed to investigate this possibility.

### 4.2 Corpus Callosum in Down syndrome

We did not detect significant differences in the volume or thickness of the corpus callosum in Ts65Dn animals (Figures 1E & 2A). This study is the first to examine the development of interhemispheric axon tracts in Down syndrome, however a previous study has examined the formation of the neocortical layers in fetal Ts65Dn animals. In E13.5 to E16.5 Ts65Dn brains, the thickness of all neocortical layers is significantly reduced as compared to euploid brains. However, this is a transient delay in neocortical layer expansion which rectified to normal values by E18.5. However, the delayed expansion of cortical layers in Ts65Dn brains might not completely rescue the connectivity and lead to abnormal brain pathology in Down syndrome [14]. A previous study also demonstrated that there was no change in the size of the corpus callosum in Ts65Dn mice at postnatal day 45 [16]. In human fetal Down syndrome brains, no study has specifically examined the corpus callosum but there are some post-mortem studies examining the cortex as a whole. Similar to Ts65Dn brains, no differences were detected in cortical thickness between fetal brains from humans with Down syndrome and controls, during late gestation [36].

### 4.3 Hippocampus in Down syndrome

The current study demonstrates that P0 Ts65Dn mice have a significant reduction in the volume of the hippocampal commissure (Figure 3C). These findings are the first to demonstrate hippocampal connectivity defects in early postnatal Ts65Dn animals. Another study has demonstrated that there is also a reduction in the size of the hippocampal commissure in Ts65Dn mice at postnatal day 45 [16]. Taken together, these studies show that hippocampal commissure deficits occur during early development and persist through adolescence.

Axons from pyramidal cells make up a large portion of the hippocampal commissure and thus the reduced hippocampal commissure volume could be due to reduced cell numbers in the hippocampus. Guidi et al. (2014) examined P2 Ts65Dn mice, and demonstrated that the number of cells is reduced in layers 2/3 of the cortex (which is the main layer whose projections form the corpus callosum) and the hippocampus. Thus, decreased numbers of cells in the hippocampus may be partially responsible for the decreased size of the hippocampal commissure; however, the number of cells is reduced in both the cortex and the hippocampus, but a defect is only present in the hippocampal commissure. This lends support to our hypothesis that there is also an axon growth defect specifically in the hippocampus (Figure 4). If reduced cell number always leads to decreased projection volume, then we should have found a reduced volume of the corpus callosum as well. Interestingly, a study that quantified numbers of only pyramidal cells in the hippocampus of P6 Ts65Dn mice found no significant change as compared to control animals [17]. Pyramidal cells are the major cell type comprising the hippocampal commissure, thus this finding also supports an axon growth deficit (Figure 4). Taken together, these data show that there is a clear time-dependent developmental component to the changes that occur in Ts65Dn mice.

Studies have shown that Ts65Dn mice exhibit cognitive deficits that depend on hippocampal function [37-40]. Taken together with previous studies, our results suggest that defects in hippocampal connectivity may contribute to impaired learning and memory in Down syndrome. The types of abnormalities that occur during the critical period of brain development, such as changes in neurogenesis and connectivity, can significantly contribute to the intellectual disability phenotype of Down syndrome. There is only one study thus far that has examined connectivity changes during embryonic development in Down syndrome mice. This study focused on thalamocortical connectivity, and found that it was defective in Ts16 Down syndrome model mice [41]. It is of significant interest to further characterize connectivity defects that occur during brain development in Down syndrome model mice, but also important to take into account that the Ts16 and Ts65Dn mouse models do not accurately replicate the genetic changes that occur in the human condition [10].

Studies on adult brains from patients with Down syndrome show decreased thickness and area of the corpus callosum [5, 6] as well as decreased hippocampal volume [5]. Taken together with the current study, this suggests that: (1) delayed maturation during brain development in Down syndrome contributes towards the pathological abnormalities found in adult Down syndrome brains, and/or (2) humans with Down syndrome undergo age-related neurodegeneration. It may be that a combination of these two processes contributes to the altered connectivity in adults with Down syndrome. There is evidence that humans with Down syndrome undergo age-related degeneration [42, 43]. For example, patients with Down syndrome aged under 6 months have longer dendritic lengths and greater dendritic branching; however as they age, dendritic arborization decreases [44]. These morphological changes may contribute to abnormal neuronal connectivity and formation of aberrant local networks. Thus, both local and interhemispheric neuronal connectivity defects may contribute to the intellectual disability phenotype of Down syndrome. Because our study demonstrates the corpus callosum is normal at P0 but the hippocampal commissure is not, it may be that the corpus callosum defects that are seen in adults with Down syndrome [5, 6] occur as a result of age-related degeneration, but the hippocampal defects are due to developmental changes. This type of knowledge is critical for us to be able to design temporally-appropriate treatments to target the various neurological phenotypes of Down syndrome.

### 4.4 Conclusions

In this study, we find that hippocampal axon length is stunted in a mouse model of Down syndrome. This is in line with our *in vivo* results that demonstrate a significant defect in hippocampal commissure connectivity in early postnatal Ts65Dn brains. These changes in connectivity may contribute to the intellectual disability phenotype of Down syndrome, however further studies are needed to investigate this possibility. Using both mouse models and iPSC-derived neurons from individuals with Down syndrome, future studies will examine the mechanisms underlying altered connectivity in Down syndrome.

## Author Contributions

SJ helped to design experiments, performed research on a daily basis, wrote the initial draft of the manuscript and edited the manuscript. CW helped to design experiments and performed research. WC helped to design experiments and edited the manuscript. KW initiated the project, designed experiments, supervised all experiments on a daily basis, and wrote the manuscript.

## Acknowledgments

We thank Dr. Guofa Liu for the kind gift of the DSCAM antibody. Ts65Dn mice are supported by NICHD contract #275201000006C-3-01.

## References

[1] G. de Graaf, F. Buckley, B.G. Skotko, Estimates of the live births, natural losses, and elective terminations with Down syndrome in the United States, Am J Med Genet A 167A (2015) 756-767.

[2] B. Schmidt-Sidor, K.E. Wisniewski, T.H. Shepard, E.A. Sersen, Brain growth in Down syndrome subjects 15 to 22 weeks of gestational age and birth to 60 months, Clin Neuropathol 9 (1990) 181-190.

[3] K.E. Wisniewski, Down-Syndrome Children Often Have Brain with Maturation Delay, Retardation of Growth, and Cortical Dysgenesis, Am J Med Genet (1990) 274-281.

[4] A. Contestabile, T. Fila, C. Ceccarelli, P. Bonasoni, L. Bonapace, D. Santini, R. Bartesaghi, E. Ciani, Cell cycle alteration and decreased cell proliferation in the hippocampal dentate gyrus and in the neocortical germinal matrix of fetuses with Down syndrome and in Ts65Dn mice, Hippocampus 17 (2007) 665-678.

[5] S.J. Teipel, M.B. Schapiro, G.E. Alexander, J.S. Krasuski, B. Horwitz, C. Hoehne, H.J. Moller, S.I. Rapoport, H. Hampel, Relation of corpus callosum and hippocampal size to age in nondemented adults with Down’s syndrome, Am J Psychiatry 160 (2003) 1870-1878.

[6] P.P. Wang, S. Doherty, J.R. Hesselink, U. Bellugi, Callosal morphology concurs with neurobehavioral and neuropathological findings in two neurodevelopmental disorders, Arch Neurol 49 (1992) 407-411.

[7] P.E. Sylvester, The anterior commissure in Down’s syndrome, J Ment Defic Res 30 (Pt 1) (1986) 19-26.

[8] M.T.S. Davisson C.; Reeves, R.H.; Irving, N.G.; Akeson, E.C.; Harris, B.S.; Bronson, R.T., Segmental trisomy as a mouse model for Down syndrome, Progress in clinical and biological research 384 (1993) 117-133.

[9] I. Das, R.H. Reeves, The use of mouse models to understand and improve cognitive deficits in Down syndrome, Dis Model Mech 4 (2011) 596-606.

[10] M. Gupta, A.R. Dhanasekaran, K.J. Gardiner, Mouse models of Down syndrome: gene content and consequences, Mamm Genome 27 (2016) 538-555.

[11] A. Duchon, M. Raveau, C. Chevalier, V. Nalesso, A.J. Sharp, Y. Herault, Identification of the translocation breakpoints in the Ts65Dn and Ts1Cje mouse lines: relevance for modeling Down syndrome, Mamm Genome 22 (2011) 674-684.

[12] P.L. Roubertoux, M. Carlier, Mouse models of cognitive disabilities in trisomy 21 (Down syndrome), Am J Med Genet C Semin Med Genet 154C (2010) 400-416.

[13] M. Dierssen, Down syndrome: the brain in trisomic mode, Nature Reviews Neuroscience 13 (2012) 844-858.

[14] L. Chakrabarti, Z. Galdzicki, T.F. Haydar, Defects in embryonic neurogenesis and initial synapse formation in the forebrain of the Ts65Dn mouse model of Down syndrome, J Neurosci 27 (2007) 11483-11495.

[15] S. Guidi, P. Bonasoni, C. Ceccarelli, D. Santini, F. Gualtieri, E. Ciani, R. Bartesaghi, Neurogenesis impairment and increased cell death reduce total neuron number in the hippocampal region of fetuses with Down syndrome, Brain Pathol 18 (2008) 180-197.

[16] S. Guidi, F. Stagni, P. Bianchi, E. Ciani, A. Giacomini, M. De Franceschi, R. Moldrich, N. Kurniawan, K. Mardon, A. Giuliani, L. Calza, R. Bartesaghi, Prenatal pharmacotherapy rescues brain development in a Down’s syndrome mouse model, Brain 137 (2014) 380-401.

[17] H.A. Lorenzi, R.H. Reeves, Hippocampal hypocellularity in the Ts65Dn mouse originates early in development, Brain Res 1104 (2006) 153-159.

[18] I. Das, J.M. Park, J.H. Shin, S.K. Jeon, H. Lorenzi, D.J. Linden, P.F. Worley, R.H. Reeves, Hedgehog agonist therapy corrects structural and cognitive deficits in a Down syndrome mouse model, Sci Transl Med 5 (2013) 201ra120.

[19] L.G. Reinholdt, Y. Ding, G.J. Gilbert, A. Czechanski, J.P. Solzak, R.J. Roper, M.T. Johnson, L.R. Donahue, C. Lutz, M.T. Davisson, Molecular characterization of the translocation breakpoints in the Down syndrome mouse model Ts65Dn, Mamm Genome 22 (2011) 685-691.

[20] S. Jain, K. Welshhans, Netrin-1 induces local translation of down syndrome cell adhesion molecule in axonal growth cones, Dev Neurobiol 76 (2016) 799-816.

[21] S. Kaech, G. Banker, Culturing hippocampal neurons, Nat Protoc 1 (2006) 2406-2415.

[22] K. Welshhans, G.J. Bassell, Netrin-1-induced local beta-actin synthesis and growth cone guidance requires zipcode binding protein 1, J Neurosci 31 (2011) 9800-9813.

[23] E.J. Hoffman, C.D. Mintz, S. Wang, D.G. McNickle, S.R. Salton, D.L. Benson, Effects of ethanol on axon outgrowth and branching in developing rat cortical neurons, Neuroscience 157 (2008) 556-565.

[24] L. Li, B.I. Hutchins, K. Kalil, Wnt5a induces simultaneous cortical axon outgrowth and repulsive axon guidance through distinct signaling mechanisms, J Neurosci 29 (2009) 5873-5883.

[25] H. Matsumoto, M. Nagashima, Shift in the function of netrin-1 from axon outgrowth to axon branching in developing cerebral cortical neurons, BMC Neurosci 18 (2017) 74.

[26] S. Gotti, E. Caricati, G. Panzica, Alterations of brain circuits in Down syndrome murine models, J Chem Neuroanat 42 (2011) 317-326.

[27] I.T. Lott, Neurological phenotypes for Down syndrome across the life span, Prog Brain Res 197 (2012) 101-121.

[28] L.J. Sosa, N.L. Postma, A. Estrada-Bernal, M. Hanna, R. Guo, J. Busciglio, K.H. Pfenninger, Dosage of amyloid precursor protein affects axonal contact guidance in Down syndrome, FASEB J 28 (2014) 195-205.

[29] S. Bahn, M. Mimmack, M. Ryan, M.A. Caldwell, E. Jauniaux, M. Starkey, C.N. Svendsen, P. Emson, Neuronal target genes of the neuron-restrictive silencer factor in neurospheres derived from fetuses with Down’s syndrome: a gene expression study, The Lancet 359 (2002) 310-315.

[30] H.Q. Huo, Z.Y. Qu, F. Yuan, L. Ma, L. Yao, M. Xu, Y. Hu, J. Ji, A. Bhattacharyya, S.C. Zhang, Y. Liu, Modeling Down Syndrome with Patient iPSCs Reveals Cellular and Migration Deficits of GABAergic Neurons, Stem Cell Reports 10 (2018) 1251-1266.

[31] K.L. Agarwala, S. Nakamura, Y. Tsutsumi, K. Yamakawa, Down syndrome cell adhesion molecule DSCAM mediates homophilic intercellular adhesion, Brain Res Mol Brain Res 79 (2000) 118-126.

[32] K.L. Agarwala, S. Ganesh, T. Suzuki, T. Akagi, K. Kaneko, K. Amano, Y. Tsutsumi, K. Yamaguchi, T. Hashikawa, K. Yamakawa, Dscam is associated with axonal and dendritic features of neuronal cells, Journal of Neuroscience Research 66 (2001) 337-346.

[33] G.M. Barlow, B. Micales, G.E. Lyons, J.R. Korenberg, Down Syndrome Cell Adhesion Molecule is conserved in mouse and highly expressed in the adult mouse brain, Cytogenet Cell Genet 94 (2001) 155-162.

[34] K. Yamakawa, Y.K. Huo, M.A. Haendel, R. Hubert, X.N. Chen, G.E. Lyons, J.R. Korenberg, DSCAM: a novel member of the immunoglobulin superfamily maps in a Down syndrome region and is involved in the development of the nervous system, Human Molecular Genetics 7 (1998) 227-237.

[35] Y. Saito, A. Oka, M. Mizuguchi, K. Motonaga, Y. Mori, L.E. Becker, K. Arima, J. Miyauchi, S. Takashima, The developmental and aging changes of Down’s syndrome cell adhesion molecule expression in normal and Down’s syndrome brains, Acta Neuropathol 100 (2000) 654-664.

[36] J.A. Golden, B.T. Hyman, Development of the superior temporal neocortex is anomalous in trisomy 21, J Neuropathol Exp Neurol 53 (1994) 513-520.

[37] M.E. Coussons-Read, L.S. Crnic, Behavioral assessment of the Ts65Dn mouse, a model for Down syndrome: altered behavior in the elevated plus maze and open field, Behav Genet 26 (1996) 7-13.

[38] R.M. Escorihuela, A. Fernandezteruel, I.F. Vallina, C. Baamonde, M.A. Lumbreras, M. Dierssen, A. Tobena, J. Florez, A Behavioral-Assessment of Ts65dn Mice - a Putative down-Syndrome Model, Neuroscience Letters 199 (1995) 143-146.

[39] M. Faizi, P.L. Bader, C. Tun, A. Encarnacion, A. Kleschevnikov, P. Belichenko, N. Saw, M. Priestley, R.W. Tsien, W.C. Mobley, M. Shamloo, Comprehensive behavioral phenotyping of Ts65Dn mouse model of Down syndrome: activation of beta1-adrenergic receptor by xamoterol as a potential cognitive enhancer, Neurobiol Dis 43 (2011) 397-413.

[40] R.H. Reeves, N.G. Irving, T.H. Moran, A. Wohn, C. Kitt, S.S. Sisodia, C. Schmidt, R.T. Bronson, M.T. Davisson, A mouse model for Down syndrome exhibits learning and behaviour deficits, Nat Genet 11 (1995) 177-184.

[41] A. Cheng, T.F. Haydar, P.J. Yarowsky, B.K. Krueger, Concurrent generation of subplate and cortical plate neurons in developing trisomy 16 mouse cortex, Dev Neurosci 26 (2004) 255-265.

[42] D.M. Holtzman, D. Santucci, J. Kilbridge, J. ChuaCouzens, D.J. Fontana, S.E. Daniels, R.M. Johnson, K. Chen, Y.L. Sun, E. Carlson, E. Alleva, C.J. Epstein, W.C. Mobley, Developmental abnormalities and age-related neurodegeneration in a mouse model of Down syndrome, P Natl Acad Sci USA 93 (1996) 13333-13338.

[43] J.P. Lockrow, A.M. Fortress, A.C. Granholm, Age-related neurodegeneration and memory loss in down syndrome, Curr Gerontol Geriatr Res 2012 (2012) 463909.

[44] L. Becker, T. Mito, S. Takashima, K. Onodera, Growth and development of the brain in Down syndrome, Prog Clin Biol Res 373 (1991) 133-152.

